# A systematic analysis of machine learning pipelines for robust antimicrobial resistance prediction

**DOI:** 10.64898/2026.06.28.734076

**Authors:** Alex Aselstyne, Enamundram Naga Karthik, Meriem El Azami, Romain Pogorelcnik, Quentin Fournier, Sarath Chandar

## Abstract

**Motivation:** Antimicrobial resistance (AMR) has been identified as a top global public health threat. Accurate AMR phenotype prediction from whole-genome sequencing data is an essential tool for accelerating clinical decision-making and mitigating resistance spread. Although many previous works have explored the use of tree-based machine learning (ML) models to predict resistance, the field lacks a systematic evaluation of the training pipeline across a variety of pathogenic species and antibiotics.

**Results:** Using nine clinically relevant species–antibiotic combinations from the NCBI antimicrobial susceptibility testing database, we present a detailed analysis of the ML pipeline and identify key factors affecting model performance and evaluation. We begin by relabelling all isolates using current CLSI minimum inhibitory concentration breakpoints to resolve inconsistencies and increase available data, resulting in up to a 19% label swap and 56% data enlargement per species– antibiotic combination. We identify several key training parameters including *k*-mer length, which can increase classification F1 scores by over 20 points compared to commonly used *k*-values, feature matrix truncation, which can induce polynomial time reductions with limited performance reduction, and ML model class. By comparing 5-fold cross-validation with evaluation on an unseen clinical dataset, we show that random cross-validation splits—often criticized as overly optimistic—can act as a strong proxy for downstream clinical performance, yielding closer F1 scores than phylogeny-aware splits in all cases. We finally present an interpretability study which shows that over 95% of *k*-mers used by our models are associated with identifiable genomic features. Our results highlight the importance of feature design, evaluation protocol, and biological analysis in genomic AMR prediction, and support tree-based models as a robust and interpretable method.

**Availability and implementation:** Python code is made freely available: https://github.com/chandar-lab/amr-pred

## 1. Introduction

Since the introduction of antibiotics in the mid-twentieth century, modern healthcare has relied on them to substantially reduce the mortality and morbidity of bacterial infections. Antimicrobial resistance (AMR), the ability of bacteria to survive antibiotic exposure, therefore represents an alarming threat to global population health (Aljeldah, 2022; WHO, 2023). The World Health Organization has identified 15 priority bacterial pathogens that have developed resistance to last-resort antibiotics (WHO, 2024) and observed increasing antibiotic resistance among 40% of tracked pathogen–antibiotic combinations between 2018 and 2023 (WHO, 2025). An estimated 1.14 million deaths in 2021 are attributed to infections from resistant bacterial strains, and a total of 4.71 million deaths are associated with these infections (Naghavi et al., 2024). A major contributor to the spread of AMR among bacterial populations is the complex and time-consuming process of determining antimicrobial susceptibility (Bharathiraja and Tamilmani, 2026; Ventola, 2015). Without efficient methods to identify the optimal antibiotic to target a given infection, last-resort antibiotics are being overused (Sturm et al., 2024), increasing selective pressure for resistant bacterial populations. Cost-effective and timely antimicrobial testing would reduce reliance on broad-spectrum antibiotics while also allowing targeted prescription, improving treatment outcomes, and reducing spending on ineffective antibiotics.

Traditionally, laboratory antimicrobial susceptibility testing (AST) has been used to determine the susceptibility profile of a bacterial isolate (Sanchez et al., 2025). This process begins with the isolation and culture of bacteria from a clinical specimen. Following this, the broth microdilution method is used to determine the lowest concentration of a single antibiotic that stops bacterial growth, known as the minimum inhibitory concentration (MIC). Although this process enables direct assessment of the phenotype of interest, it suffers from several limitations, including reliance on specialized facilities and trained personnel (Boolchandani et al., 2019) as well as lengthy bacterial culture growth. As an efficient alternative to wet-lab testing, computational methods have been proposed to predict phenotypic resistance from bacterial genomes (Behling et al., 2023; Ali et al., 2023). These methods fall into two broad categories: rule-based and machine learning (Hu et al., 2024). Rule-based methods, such as ResFinder (Zankari et al., 2012), use databases of known genetic markers to accurately predict resistance. The primary limitation of this approach is that resistance markers must already be known. Machine learning (ML) methods allow learning directly from data, without relying on hand-crafted rules. This makes the ML approach more versatile and better suited to understudied or novel species and drugs, while sacrificing curated, evidence-based priors.

Several studies have explored ML-based approaches for resistance phenotype prediction. Recent surveys of the literature (Lv and Wang, 2024; de la Lastra et al., 2024) highlight the prominence of tree-based models that operate on whole-genome sequences. Although neural network-based approaches have recently gained popularity across a variety of fields, their requirement for large-scale training datasets limits their utility for phenotype prediction (Grinsztajn et al., 2022). However, many works that propose ML models for AMR prediction fail to explore the breadth of possible model settings. For instance, models are often only optimized for and evaluated on a single species or antibiotic (Khaledi et al., 2020; Nguyen et al., 2019; Tolan et al., 2025), hindering the generalization across pathogens. Evaluation is also not consistently performed on both phylogeny-aware and random cross-validation folds (Aytan-Aktug et al., 2020; Ren et al., 2021), which are relevant to distinct clinical settings. Finally, studies that represent the genome with *k-mers* often fail to evaluate the impact of their length (Wittenbach et al., 2026; Drouin et al., 2019), which fixes the genomic representation.

In order to address these identified gaps, *we systematically assess ML resistance phenotype classifiers across nine species– antibiotic combinations and three evaluation protocols to identify key methodological choices*. Although previous studies have conducted large-scale benchmarking of AMR phenotype prediction models (Hu et al., 2024), they have taken an aggregate, high-level approach to their evaluation. Our primary aim is to determine which aspects of the representation, training, and evaluation pipeline are most important to consider when creating new models for AMR prediction. Additionally, we aim to verify existing assumptions in the field through analysis of cross-validation settings and interpretability analysis. Our primary contributions are:

- We present a thorough evaluation of ML models for AMR prediction across nine clinically relevant and diverse species–antibiotic combinations using curated and relabelled data.
- We identify genomic representation parameters—primarily *k*-mer length and matrix dimensionality—and ML model selection as major determinants of predictive accuracy.
- We demonstrate the relationship between model performance and the underlying biology by comparing random and phylogeny-aware folds with an external clinical dataset and by showing that important *k*-mers correspond to known resistance-associated biomarkers.

## 2. Materials and methods

### 2.1. Data selection and preprocessing

Our study spans nine combinations of species and antibiotics: *A. baumannii*–ciprofloxacin, *K. pneumoniae*–tobramycin, *S. aureus*–erythromycin, *A. baumannii*–ceftazidime, *E. coli*– cefotaxime, *K. pneumoniae*–meropenem, *P. aeruginosa*– aztreonam, *E. coli*–cefepime, and *E. coli*–imipenem. We select these combinations to cover diverse pathogenic species, antibiotic classes, resistance mechanisms, data quantities, and class imbalances. To avoid biasing our results, combination selection occurred prior to experimentation.

We begin by downloading the corresponding nine datasets from the Antimicrobial Susceptibility Testing (AST) browser of the National Center for Biotechnology Information (NCBI) BioSample database^1^ (accessed on July 22, 2025). The datasets include raw genome sequences along with associated metadata, including resistance phenotypes, MIC values, and sample origin information. Genomes marked with either resistant, susceptible, or not defined phenotypes are retained, resulting in a total of 12,741 genomes across all nine datasets. To ensure quality, consistency, and maximal data utilization, we revise all resistance phenotypes using current MIC breakpoints from the Clinical and Laboratory Standards Institute (CLSI) M100 (35th edition) (Lewis II, 2025). Given that the labels provided in the NCBI AST database may have been assigned under different testing standards, this ensures consistency in MIC to phenotype mapping. This also ensures that phenotype predictions made by our models correspond to current laboratory recommendations. As additional preprocessing steps, any samples with missing MIC values are removed and all *Shigella* samples are removed from the *E. coli* dataset. Table 1 breaks down the frequency distribution of each dataset after preprocessing.

**Table 1.** Dataset composition for each combination of pathogen and antimicrobial studied. The number of susceptible and resistant samples are determined after MIC reinterpretation using CLSI breakpoints.

| Pathogen | Antimicrobial | *n* | Susceptible Samples | Resistant Samples |
| --- | --- | --- | --- | --- |
| *Acinetobacter baumannii* | Ceftazidime | 1,170 | 284 (24%) | 886 (76%) |
|  | Ciprofloxacin | 1,482 | 284 (19%) | 1,198 (81%) |
| *Escherichia coli* | Cefepime | 3,031 | 2,624 (87%) | 407 (13%) |
|  | Cefotaxime | 751 | 182 (24%) | 569 (76%) |
|  | Imipenem | 4,199 | 4,153 (99%) | 46 (1%) |
| *Klebsiella pneumoniae* | Meropenem | 839 | 372 (44%) | 467 (56%) |
|  | Tobramycin | 525 | 202 (38%) | 323 (62%) |
| *Pseudomonas aeruginosa* | Aztreonam | 144 | 54 (38%) | 90 (62%) |
| *Staphylococcus aureus* | Erythromycin | 600 | 305 (51%) | 295 (49%) |

### 2.2. Cross-validation fold generation

To match the evaluation standards of previous studies, we evaluate using two types of dataset splits for our five-fold cross-validation: random and phylogeny-aware. Random splits generate five disjoint folds (groups) of the data at random without accounting for the evolutionary relationships between isolates. As a result, randomly-generated cross-validation folds are believed to fail to capture the genomic diversity seen in some clinical and environmental settings (Yu et al., 2025). In contrast, phylogeny-aware folds take into account the distance between the genomes of isolates, creating diverse training and testing data (Hu et al., 2024; Lüftinger et al., 2021). Random folds are generated using stratified five-fold cross-validation (through the StratifiedKFold method available in the scikit-learn Python package) with a fixed random seed to ensure reproducibility. To create phylogeny-aware folds, a dendrogram is constructed using a distance matrix relating all samples in each species–antibiotic dataset. The dendrogram is then partitioned into five groups of approximately the same size, which we define as being within five isolates, with maximum dissimilarity between them. This results in five disjoint and genetically dissimilar folds. We compare evaluation on random and phylogeny-aware cross-validation folds to quantify the performance delta between them.

In the generation of our phylogeny-aware folds, we test three genomic distance tools: MUMmer3 (Kurtz et al., 2004), Mash (Ondov et al., 2016), and Dashing (Baker and Langmead, 2018). We use MUMmer3 to calculate average nucleotide identity (ANI) as our ground truth distance metric, while Mash and Dashing both output Mash distance, a strong proxy for ANI. We test all three tools to determine an adequate balance of speed and accuracy, varying the sketch size parameter for both Mash and Dashing. Following our experiments, we determined that Dashing with a sketch size of 2^10^ provided the best balance of speed and accuracy. Related results are provided in Section A in the Supplementary Notes.

### 2.3. Genomic feature representation

A common way to represent long genome strings for input to an ML model is using *k*-mer matrices (Drouin et al., 2014; Khaledi et al., 2020). To construct our matrix, we begin by extracting all *k*-mers (overlapping subsequences of size k) present in the training data, with reverse compliments canonicalized. Then, for each isolate, we count the number of occurrences of each *k*-mer in the given genome. These counts are used to construct a feature matrix where rows correspond to isolates and columns to *k*-mers. While *k*-mers of sizes 21–31 are typically used for balancing computational efficiency and optimal sequencing coverage (Jenike et al., 2025), there are few meta-analyses studying the impact of *k*-mers of different sizes for resistance phenotype prediction (Chikhi and Medvedev, 2013). Therefore, we systematically evaluate *k* = *{*3, 17, 31, 45*}* to observe the impact of *k*-mer length on predictive performance across species–antibiotic combinations.

As the total number of *k*-mers grows exponentially with *k*, fitting the feature matrix into memory becomes a significant computational challenge. As a basic reduction technique, we always remove *k*-mers that appear fewer than twice across the entire dataset, as exceedingly rare, non-repetitive *k*-mers are unlikely to have discriminative power. We then experiment with two methods to more aggressively trim the matrix columns. First, we perform matrix column truncation following frequency column ordering, which involves re-ordering the columns from the most prevalent to the least prevalent *k*-mers and then trimming low-prevalence columns. We evaluate a range of matrix sizes containing the most frequently occurring 10,000 to 10 million *k*-mers. Our second, more sophisticated dimensionality reduction technique is singular value decomposition (SVD). Our aim with SVD-projection is to distill the information contained in the complete *k*-mer matrix to a significantly reduced number of features. This is an attractive option, as it enables significantly lower computational load for both training and inference while maintaining the majority of the original variability. A drawback, however, is the loss of interpretability present when using the true *k*-mers. We select SVD over principal components analysis to avoid the memory overhead of centering sparse matrices. At training time, we first perform SVD on the *k*-mer matrix and truncate the resulting singular value matrix to only the desired number of components. To compensate for the lack of centering, all component scores were mean-centered after projection. At test time, the testing data are projected onto the components learned from the training data to maintain alignment.

### 2.4. Model selection and tuning

We evaluate two model types across the two matrix reduction techniques outlined in subsection 2.3: logistic regression and XGBoost with truncated inputs, and XGBoost with SVD-transformed inputs. We select these models to cover a range of ML methods, varying in complexity. Each model is tuned using a Bayesian hyperparameter search (Bergstra et al., 2011) over 25 rounds. Compared to simple grid or random search strategies, this modern approach uses information from previous rounds to guide the selection of hyperparameters. At each iteration of the cross-validation (described in subsection 2.2), one-fifth of the training data is randomly selected as a validation subset for hyperparameter tuning. The remaining four-fifths is used to train the model with each of the tested hyperparameter settings. Our ML model hyperparameter tuning always takes place on a randomly selected validation set, regardless of the CV split method, and the same validation/training sets are used for each round of tuning within a given fold. After the optimal hyperparameters are found on the tuning subset, the model is trained with these hyperparameters on the complete training fold from the outer cross-validation (Figure 1B). The hyperparameters tuned and the ranges for each are provided in Section D in the Supplementary Notes.

**Figure 1.**
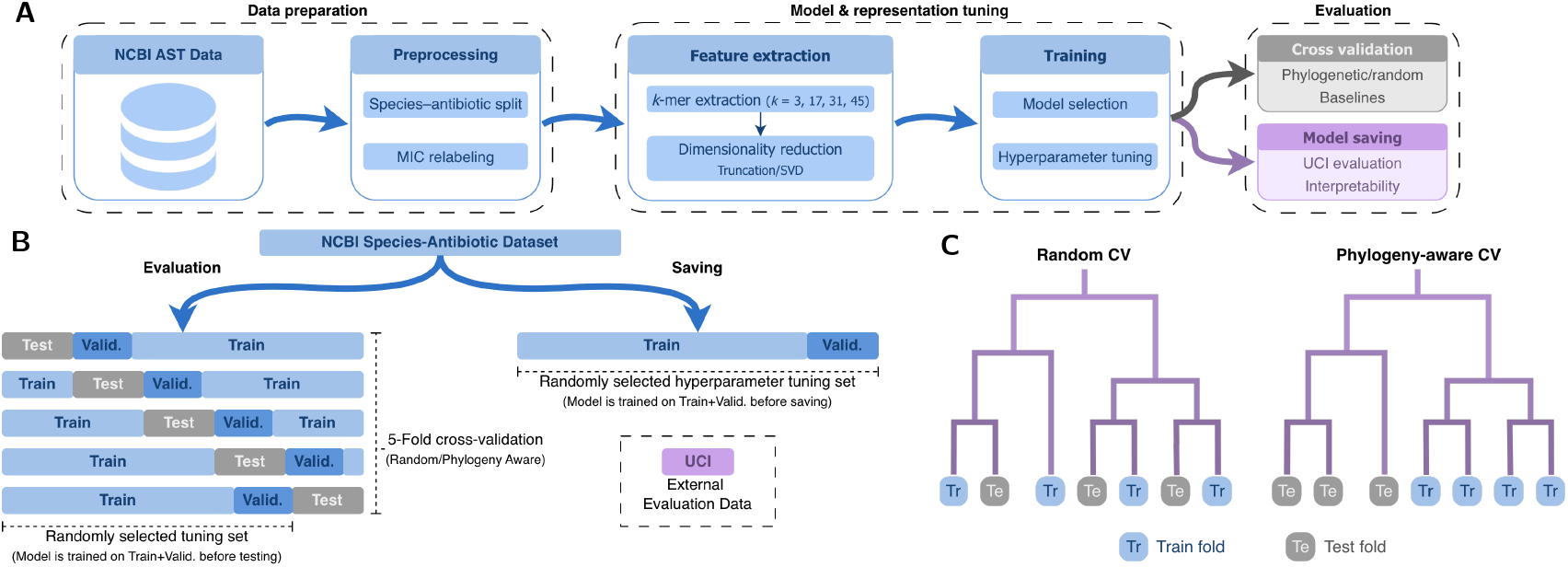
**A:** Overview of our evaluation pipeline. Results are grouped into data preparation, model & representation tuning, and finally evaluation. **B:** Diagram of data partitioning for cross-validation and model saving. When performing cross-validation, test sets are determined beforehand using either random or phylogeny-aware folds. A validation (valid.) set is always randomly selected from the remainder of the data for hyperparameter tuning. When saving models, only the random validation set is necessary. After hyperparameter tuning, models are always trained on the combined train/validation data before testing or saving. **C:** Trivial dendrogram illustrating the difference in construction of random and phylogeny-aware folds. For the latter, we construct the phylogenetic tree and partition it into 5 groups, each used as a testing fold.

When training models to be saved for external data inference and interpretability analysis, cross-validation is not used. Instead, the entire training dataset for the target species– antibiotic combination is randomly split into training (80%) and validation (20%). 25 rounds of hyperparameter tuning are performed on these splits before the final model is retrained on all available training data using the optimal hyperparameters and saved.

### 2.5. Model evaluation

When evaluating the impact of model parameters and representations, we primarily consider CV performance using NCBI data. To assess classification performance in this setting, we compute the macro F1 score across folds for each species– antibiotic combination. Although we only report macro F1 scores in the main paper, balanced accuracy, major error rates, and very major error rates were also recorded (Section J in the Supplementary Notes). We compare our method to two baseline methods: ResFinder (Zankari et al., 2012) and Aytan-Aktug (Aytan-Aktug et al., 2020). We select these baselines based on the results of Hu et al. (2024) to cover the three primary model types — rule-based, deep learning, and our own traditional machine learning. ResFinder is a rule-based approach that uses catalogues of known genetic resistance markers. Although Aytan-Aktug is an ML method, it employs a neural network instead of a tree-based approach. Additionally, we evaluate our models on an external clinical dataset (Wittenbach et al., 2026) to test performance on unseen data. We refer to these data as the *UCI dataset*, and a description of dataset composition is available in Section F in the Supplementary Notes. As we do not apply any additional preprocessing to the dataset, all details of data processing can be found in Wittenbach et al. (2026).

As part of our evaluation, we perform an interpretability analysis to determine whether the *k*-mers selected by our models align with known resistance biomarkers. Important *k*-mers are determined using SHapley Additive exPlanation (SHAP) values (Lundberg and Lee, 2017). SHAP values are selected because they provide a consistent feature-attribution framework across model types. We extract *k*-mers using the shap Python library from the final XGBoost models trained on all available NCBI data for each species–antibiotic combination. For each model, we annotate all *k*-mers that have a SHAP value of at least one-tenth of the single most important *k*-mer to target our analysis at only relevant *k*-mers. The selected *k*-mers are annotated using NCBI RefSeq genomes (Goldfarb et al., 2025). Each *k*-mer is first aligned using Burrows-Wheeler alignment (Li and Durbin, 2009) with zero-mismatch exact matching on a random selection of up to 1,000 RefSeq assemblies. Next, the alignments are intersected with the matched NCBI RefSeq annotation files (GFF) for each assembly using BEDTools (Quinlan and Hall, 2010). Finally, annotations are aggregated across assemblies, and the most frequently occurring annotation is assigned to each *k*-mer.

## 3. Results

To avoid the computational cost of a full grid search over all model and representational parameters, we ran our experiments in a greedy manner, testing only one variable at a time and utilizing the best values in subsequent experiments. Except where otherwise noted, reported metrics are averaged across all five phylogeny-aware cross-validation folds.

### 3.1. Data preparation

We began our analysis by curating quality datasets for each species–antibiotic combination of interest. Our most consequential form of preprocessing, relabelling isolates using current laboratory guidelines, resulted in an impactful change in the composition of the datasets (Figure 2). This is especially true for *E. coli*–imipenem, where the total amount of labelled samples for training increased by 1,520 (Section B in the Supplementary Notes) by leveraging isolates that were not previously labelled. Perhaps to greater impact, a number of species–antibiotic combinations also had isolates that switched phenotypes. The number of resistant samples for *K. pneumoniae*–meropenem, for example, increased 19.1% when 75 previously susceptible isolates flipped to resistant. This reflects the evolving nature of AMR as breakpoint changes occur in response to new data, the emergence of new resistance mechanisms, regulatory change, or the evolution of our understanding of resistance. Although fewer than 1% of isolates differed from their original labels after reinterpretation for four of nine combinations, it is clear from those remaining that relabelling is a critical preprocessing step to maximize the currency, consistency, and utilization of available data.

**Figure 2.**
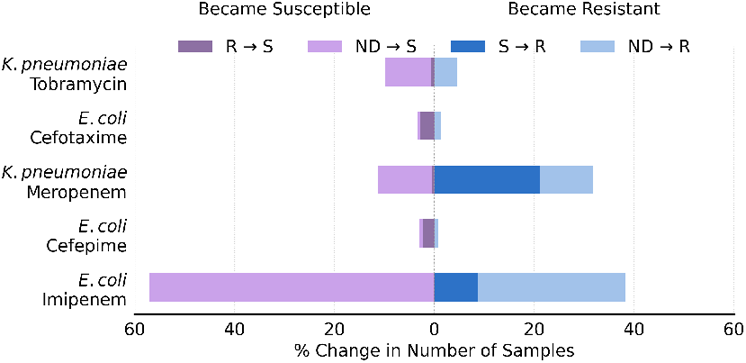
Percent change in number of samples per combination for each label. Percentages are calculated relative to the original number of isolates classified with the final label. Species for which *<* 1% of samples changed are omitted. Lighter colours correspond to added data, and darker to swapped labels.

After reinterpretation, our selected combinations cover a variety of possible data availability settings (Table 1). *K. pneumoniae*–meropenem and *S. aureus*–erythromycin represent comparatively easy combinations for learning; both datasets are balanced between label classes, with susceptible/resistant proportions of 44%/56% and 51%/49%, and contain a substantial number of isolates (839 and 600, respectively). In contrast, *E. coli*–imipenem and *P. aeruginosa*–aztreonam present difficulties for model training, with extremely imbalanced classes (99%/1%) and limited data (144 isolates), respectively. We evaluate across these diverse settings to identify common model failure modes and broader trends in the landscape of bacterial pathogens. The first stage of this evaluation is tuning, where we select the optimal values of training parameters.

### 3.2. Model and representation tuning

A critically overlooked aspect of model training throughout the literature is the impact of the *k*-mer length, the basis of genomic representation. We found that tuning the *k*-value for each species–antibiotic combination resulted in considerable macro F1 gains over the universal application of any one *k*. Of the four *k*-mer lengths tested, *k* = *{*3, 17, 31, 45*}*, each was found to result in the highest macro F1 score for at least one combination of species and antimicrobial. Of the nine combinations tested, six performed best at *k* = 17. The remaining three combinations (Figure 3A) performed best at differing *k*-values, with *A. baumannii*–ceftazidime at *k* = 31, *E. coli*–imipenem at *k* = 45 and *P. aeruginosa*–aztreonam at *k* = 3.Trends in performance characteristics across *k*-mer lengths were also not observed to be consistent. Some combinations, such as *S. aureus*–erythromycin, have very sharp distributions, while others like *K. pneumoniae*–tobramycin are much flatter. Taken together, these inconsistencies suggest that it is best to optimize *k* for each combination to maximize accuracy.

**Figure 3.**
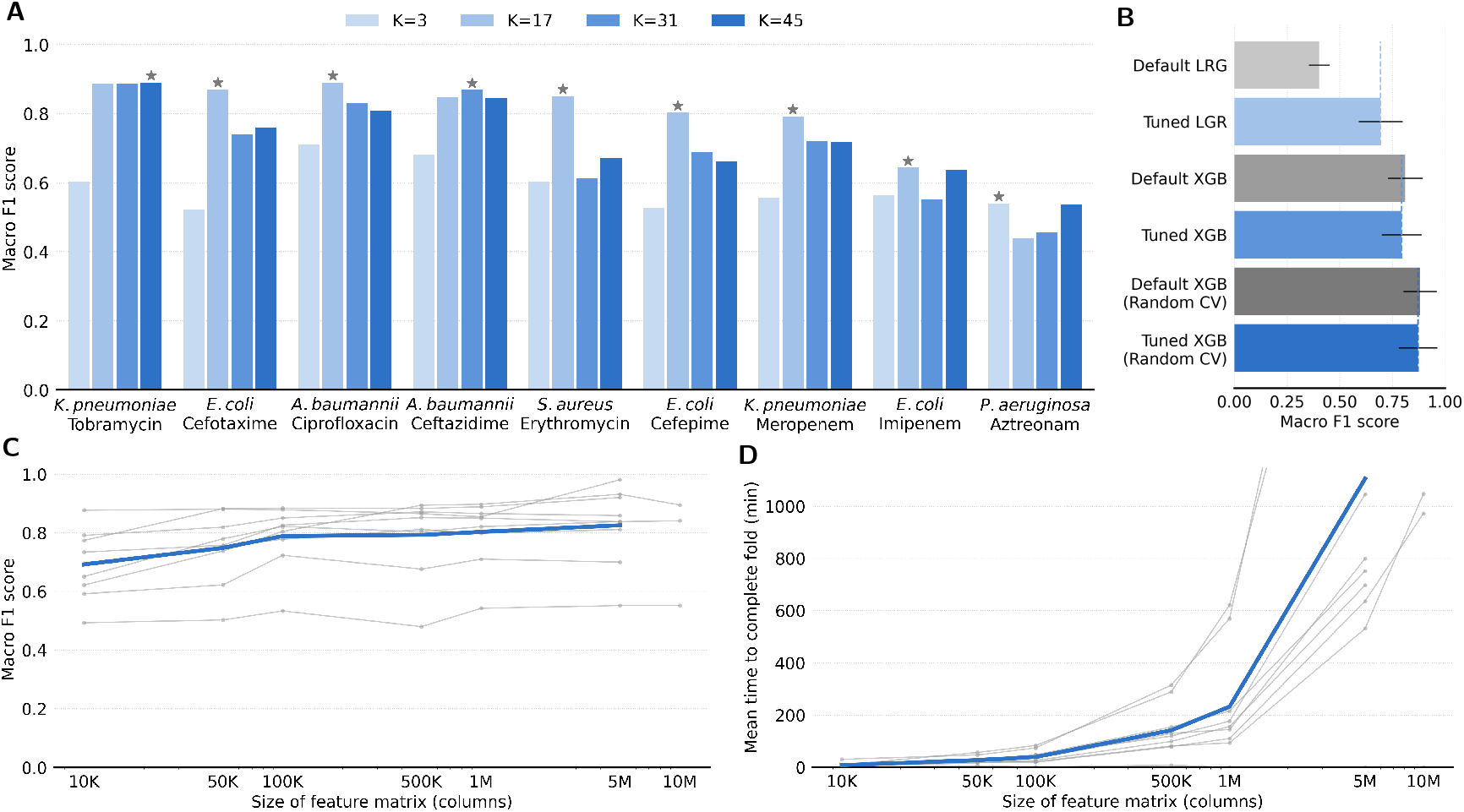
**A:** Impact of *k*-mer length on model performance for all tested combinations. The optimal *k*-value for each is denoted with a star. **B:** Macro F1 scores on held out test sets from 5-fold CV for default and tuned hyperparameter settings. Logistic regression (LGR) sees a significant improvement, while XGBoost (XGB) does not. **C:** Model performance across feature matrix sizes (log scaled), pruned using truncation. Average of all models is shown in blue. Performance increases slowly with additional *k*-mers. **D:** Mean minutes to complete a training fold, with hyperparameter tuning, at each feature matrix size. Time complexity increases polynomially with dimensionality.

**Figure 4.**
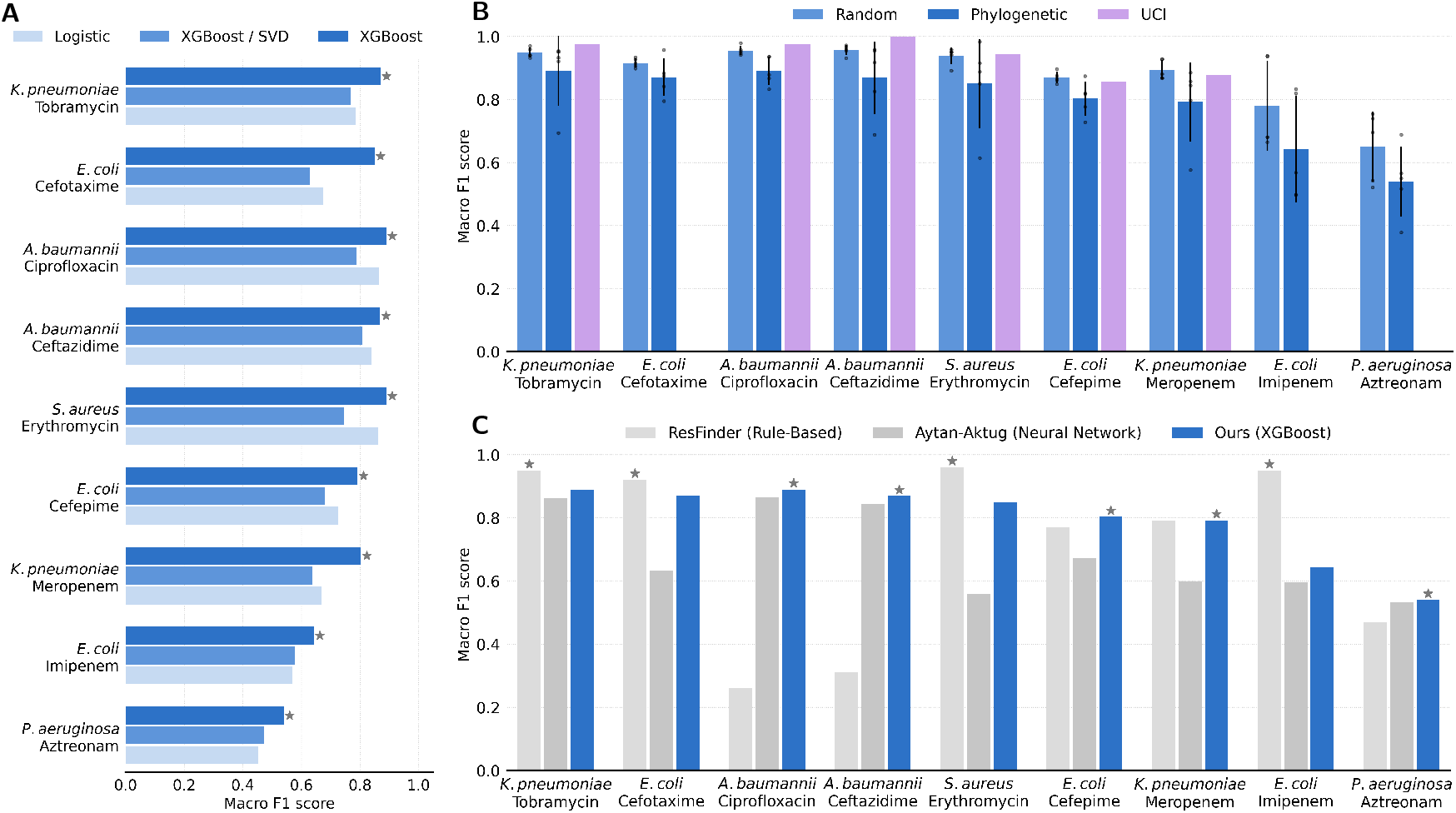
**A:** Performance of logistic regression, XGBoost with SVD matrices, and XGBoost with regular *k*-mer matrices. **B:** Model evaluation using random folds, phylogeny-aware folds, and the UCI dataset. Phylogeny-aware folds are consistently more difficult than the other two evaluation settings. **C:** Performance comparison with rule-based (ResFinder) and neural network (Aytan-Aktug) baselines across studied combinations. ResFinder achieves the highest macro F1 when rules are available, and our tree-based method performs better than the neural network baseline.

Another important facet of model tuning is feature matrix size, as it is the primary computational bottleneck. We found that *k*-mer matrices with more columns tended to lead to higher F1 scores, indicating that very uncommon *k*-mers can still be critical to model decisions. An exception to this trend is seen in *P. aeruginosa*–aztreonam, the lowest grey line in Figure 3C, as *k* = 3 only allows for a maximum of 32 columns. Contrary to our expectations, F1 scores occasionally decrease when the size of the *k*-mer matrix is increased. Because matrices with fewer columns are subsets of larger ones, models trained on the larger matrices can recover the same informative *k*-mers and thus achieve at least comparable performance. We attribute the observed performance drops to overfitting on rare *k*-mers.

Although average macro F1 steadily increases with larger feature matrices, the mean time required to complete one training fold follows a power law (Figure 3D). This difference in compute requirements is present during both training and testing and, as such, is an important consideration when designing the model for downstream applications. Although larger *k*-mer matrices increase computational requirements substantially, they provide only small gains in predictive performance. Due to this highly unfavourable tradeoff between predictive accuracy and computational cost, we conduct the remainder of our experiments using 1M–column *k*-mer matrices.

The final stage of model optimization is hyperparameter tuning. To assess its impact, we compared the macro F1 scores of models optimized using 25 rounds of Bayesian hyperparameter search with those of models trained on the same folds using the default library hyperparameters. We found that hyperparameter tuning was very important for logistic regression models, resulting in a 72% increase in macro F1. For XGBoost models, however, we found that tuning made no significant impact to macro F1 scores. A Wilcoxon signed-rank test found this to be true for both phylogeny-aware folds (*W* =16.0, *p*=0.496, *n*=9) and random (*W* =7.0, *p*=0.074, *n*=9), when testing across all nine species–antibiotic combinations. Visual comparison does not reveal any specific trends across folds or species (Section I in the Supplementary Notes). These observations indicate that either the default hyperparameters for XGBoost are already approximately optimal within our search space, or XGBoost is insensitive to hyperparameter adjustments for this problem. Our results suggest that for high-performing models, hyperparameter tuning is not as important as *k*-mer length or representation dimensionality tuning.

### 3.3. Model evaluation

Our primary evaluation strategy consisted of two cross-validation settings. In all cases, evaluating with phylogeny-aware folds resulted in lower macro F1 scores than random splits. This supports the hypothesis that phylogeny-aware folds create more difficult evaluation environments. However, an important observation arose when comparing cross-validation results to performance on the UCI dataset. Since the UCI dataset is composed of samples taken from clinical settings in the United States, bacterial sources are similar to those of the NCBI AST dataset. Phylogeny-aware cross-validation splits are often purported to be more similar to real-world application settings (Roberts et al., 2017; Lüftinger et al., 2021), but our results did not support this theory. This leads us to conclude that random cross-validation splits are an appropriate measure of model performance when the target downstream deployment environment is closely related to the training environment. We therefore recommend that both random and phylogeny-aware evaluations be reported when proposing new methodologies, as they can be indicative of different clinical conditions.

Next, we evaluated the performance of matrices transformed using the SVD. We compared three model configurations: XGBoost and logistic regression using *k*-mer matrices with 1M features, and XGBoost using a 163-component SVD-reduced representation (Section H in the Supplementary Notes). We included logistic regression as a simpler baseline model type to compare with the other models. Of the three approaches, XGBoost with SVD-reduced matrices achieved the lowest F1 scores in the majority of cases. The two cases where the SVD-reduced representation outperformed logistic regression yielded F1 scores close to 0.5, indicating limited practical relevance. This suggests either that the components do not capture the genomic variation relevant for phenotypic resistance or that a greater number of components are required to adequately model the underlying variance. The best performing configuration in all tested cases was XGBoost with untransformed *k*-mer matrices. A notable advantage of SVD matrices is the significant reduction in both training and testing times, even accounting for SVD learning and projection (Section E in the Supplementary Notes). For this reason, reduced-size feature matrices remain an interesting future research direction.

Finally, we compared our model to existing baseline methods to assess performance in varying prediction strategies. Generally, we found that ResFinder, a rule-based approach, achieved higher macro F1 scores than the other methods when biomarker catalogues are available for a given combination. This was especially true in cases where learning was challenging, such as the extremely imbalanced *E. coli*–imipenem dataset. When specific rules are not available and ResFinder has to fall back on general rules, however, it performed significantly worse than both the neural network and tree-based models. This highlights the key shortcoming of rule-based models: handcrafted features must be determined ahead of time. This limits their applicability in emerging drug or understudied pathogen settings. The Aytan-Aktug neural network baseline underperformed compared to our tree-based approach in our low-data setting.

### 3.4. Interpretability

Biological interpretability plays an essential role in model evaluation, as it roots predictions in empirical evidence and strengthens trust in model outputs. We chose to analyze *k*-mers that are highly important to each model, specifically those with at least one tenth the SHAP value of the most important *k*-mer for each species–antibiotic combination. The number of *k*-mers analyzed via this method was not correlated with the classification F1 scores (*r* = *−*0.294, *p* = 0.480), highlighting that some resistance phenotypes can be accurately predicted from a small number of strong biomarkers, while others are distributed across a broader set of genomic signals. When aligning each of these *k*-mers with NCBI RefSeq genomes, the associated genomic sequences were annotated with one of five categories: known resistance marker, protein coding, transfer RNA (tRNA), ribosomal RNA (rRNA), or insertion sequence (IS) element. All categories can be related to AMR (Elbaiomy et al., 2025; Razavi et al., 2020; Babosan et al., 2022). The categorical distribution was found to vary substantially between different combinations (Figure 5), highlighting the variety of complexities among resistance mechanisms. Only 4.27% of the *k*-mers analyzed were unable to be annotated, demonstrating biologically grounded predictions.

**Figure 5.**
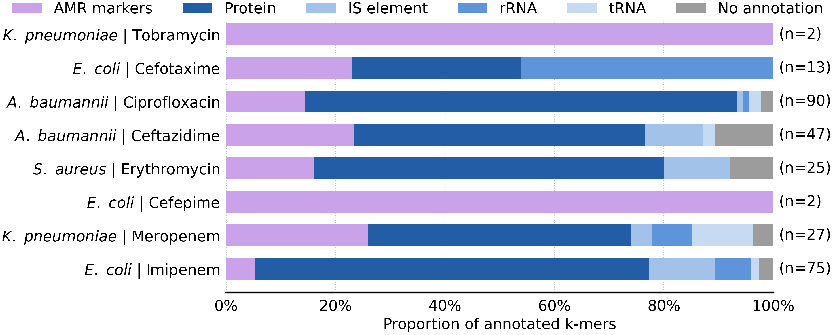
Summary of the interpretability study. Bars show proportions of all *k*-mers considered for annotation that were matched to each genetic element type. The total number of considered *k*-mers for each combination is included on the right.

Several species–antibiotic combinations demonstrated that our models rely on biologically supported resistance-associated signals. In the case of *K. pneumoniae*–tobramycin, both *k*-mers identified as important mapped to *aac(6’)-Ib*, a known aminoglycoside resistance gene, showing models can identify canonical resistance determinants. Beyond direct resistance genes, the model also captured genomic context associated with resistance. For *K. pneumoniae*–meropenem, one highly ranked *k*-mer mapped to an insertion sequence between *aac(6’)* and *blaKPC*, indicating the model may leverage mobile genetic elements linked to resistance dispersal. Finally, some combinations appeared to rely on broader cellular signals. In *E. coli*–cefotaxime, multiple highly important *k*-mers mapped to 16S rRNA regions, which are closely tied to ribosomal function. Together, these findings suggest that the model captures both established resistance biomarkers and wider genomic patterns associated with resistant phenotypes.

## 4. Discussion

Here, we present a thorough evaluation of ML models for AMR prediction. This includes evaluation of multiple model types across nine species–antibiotic combinations, using properly annotated curated data. We show that many trends in model performance are consistent across combinations, though not all are. XGBoost tree-based models are consistently better than logistic regression models, for example, but the choice between rule-based or ML predictors depends on the amount and balance of data available for a given combination, the quality of existing rules, and the type of ML model being considered. Additionally, relabelling data is a critical step for ML pipelines, as more data is beneficial to model predictions.

We also show that the process of tuning the model and its genomic representation can lead to significant performance gains. A key hyperparameter of interest is *k*-mer length, which together with model type can make a significant difference in classification scores. We found that XGBoost models on truncated *k*-mer matrices were the best model choice tested and that truncation is a very unbalanced tradeoff between macro F1 and compute, suggesting that a good representational choice is *k*-mer matrices of a tuned length truncated to the one million most common features. Additionally, we determined that depending on the evaluation setting, ML model hyperparameter tuning can vary from unimpactful to extremely beneficial.

Finally, we show that model performance is strongly related to the underlying biology through our interpretability analysis and validation settings. Our interpretability study found that virtually all (95%) of *k*-mers could be mapped to a reference genetic element, and our cross-validation pipeline showed that phylogeny-aware folds underestimated performance on clinical data. Taken together, these findings illustrate that our ML models are sensitive to the evolutionary relationships between bacteria and can identify key genomic regions affecting resistance. This supports the potential of ML models as robust methods for AMR phenotype prediction, and we recommend that future practitioners evaluate models using both random and phylogeny-aware cross-validation folds.

A limitation of our work is the focus on a single-model-per-combination setting, while a number of models have also been proposed to predict across multiple species or antibiotics (Hu et al., 2024). The field would benefit from future work performing similar studies in the multi-species and multi-antibiotic settings. Additionally, a strong future direction is the exploration of models that predict MIC values directly instead of binary labels. This regression framing provides more granular predictions and allows resistance phenotypes to be adjusted to new breakpoints at any time. Finally, the recent widespread application of foundation models to a variety of domains has inspired us to question their applicability for AMR phenotype prediction. A number of genomic foundation models (Nguyen et al., 2024; Boshar et al., 2025) have been proposed, and their broad understanding of the genomic language may enable them to encode resistance signals. We position our work as a strong baseline to compare such methods against.

## Author contributions

A.A., E.N.K., M.E.A., and Q.F. conceived the experiments; A.A. and E.N.K. conducted the experiments; A.A., E.N.K., M.E.A., and Q.F. analyzed the results; R.P. and M.E.A. conducted and analyzed the interpretability study; A.A. wrote the manuscript; all authors edited the manuscript; Q.F. and S.C. supervised.

## Competing interests

M.E.A. and R.P. are employees of bioMérieux SA, a biotechnology company. The authors declare no other competing interests.

## Acknowledgments

We acknowledge Arthur Bernard and Farah Ben Slama from bioMérieux SA, as well as Lola Le Breton from the Chandar Lab at Mila for their insightful discussions and feedback. This research was supported by bioMérieux SA. A.A. is supported by the Natural Sciences and Engineering Research Council of Canada (NSERC). S.C. is supported by the Canada CIFAR AI Chairs program, the Canada Research Chair in Lifelong Machine Learning, and the NSERC Discovery Grant. This research was enabled by compute resources provided by Mila (mila.quebec) and the Digital Research Alliance of Canada (alliancecan.ca).

## Footnotes

1 https://www.ncbi.nlm.nih.gov/pathogens/ast

## Notes

https://github.com/chandar-lab/amr-pred

### Competing Interest Statement

M.E.A. and R.P. are employees of bioMerieux SA, a biotechnology company. The authors declare no other competing interests.

