## Supplementary Notes for "A systematic analysis of machine learning pipelines for robust antimicrobial resistance prediction"

### Supplementary Material

#### A. Distance matrix generation

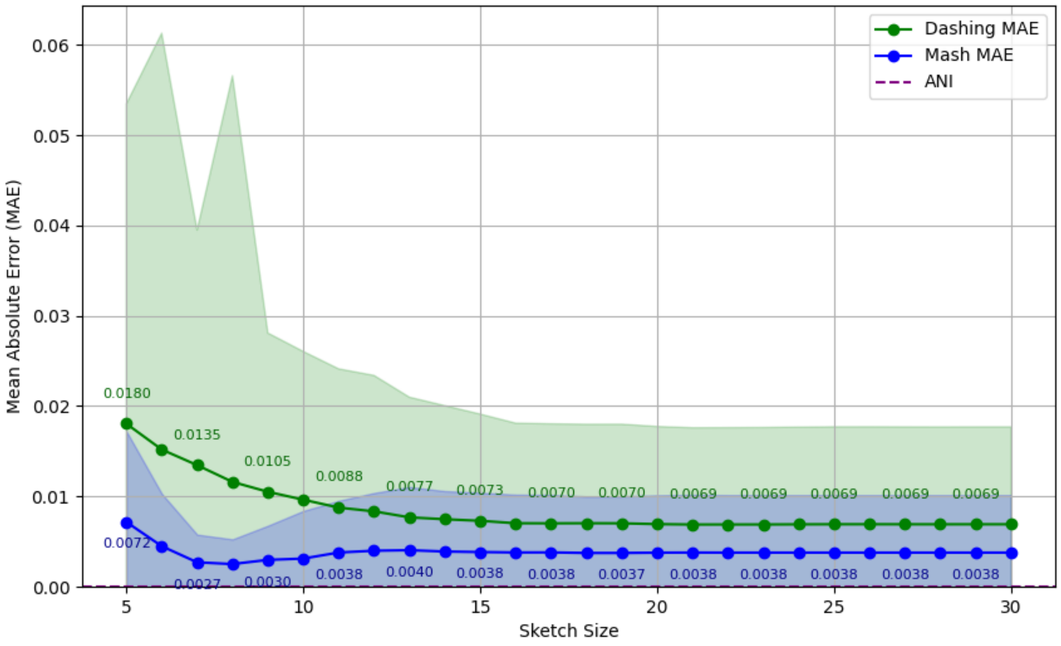


**Figure** 6: Comparison of Mean Absolute Error (MAE) of the Mash and Dashing tools at varying sketch sizes (as powers of 2), compared to ANI distance from MUMmer3. Values are averaged across 100 *A. baumannii* samples, and shaded areas represent standard deviation across samples

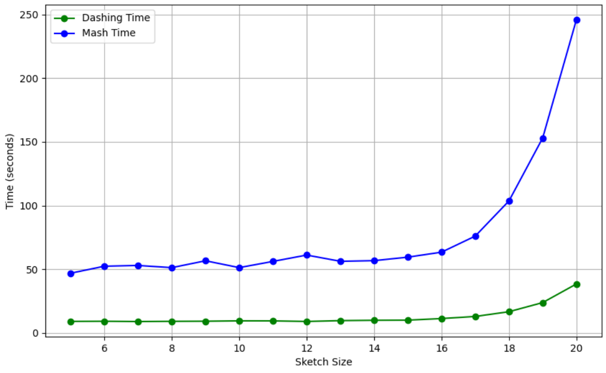


**Figure** 7: Comparison of time required to complete a full pairwise distance calculation for 100 *A. baumannii* isolates for sketch sizes from five to 20 for both Mash and Dashing.

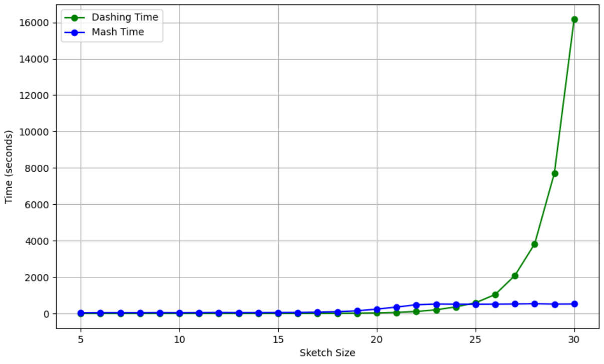


**Figure** 8: Extended version of Figure 7 that shows performance at sketch sizes above $2^{20}$, up to $2^{30}$.

In this phase of our experiments, our aim was to find the optimal tool and sketch size to generate accurate distance matrices at scale. These matrices contain pairwise genomic distances between all samples for a given species–antibiotic combination and are necessary to construct the phylogenetic tree. We elected to test Mash, Dashing, and MUMmer3. MUMmer3 computes the exact average nucleotide identity (ANI) between samples, a computationally expensive distance metric due to sequence alignment. Mash and Dashing (the more recent tool) create *k*-mer based sketches of the genomes, and use those to compute Mash distance.

Our analysis of Figures 6, 7 and 8 led us to conclude that Dashing provides a reasonable proxy that is significantly faster than Mash. For our final distance matrices, we used a sketch size of $2^{10}$, as this fell under our MAE threshold of 0.01.

#### B. Dataset reinterpretation counts

Table 2: Counts of samples changed per label after reinterpretation with CLSI MIC breakpoints.

| Species | Antibiotic | S→R | R→S | ND→R | ND→S |
|---|---|---|---|---|---|
| *A. baumannii* | Ceftazidime | - | - | 1 | - |
| | Ciprofloxacin | - | - | 1 | - |
| *E. coli* | Cefepime | - | 57 | 4 | 18 |
| | Cefotaxime | - | 5 | 8 | 1 |
| | Imipenem | 3 | 1 | 10 | 1510 |
| *K. pneumoniae* | Meropenem | 75 | 2 | 38 | 43 |
| | Tobramycin | - | 1 | 14 | 17 |
| *P. aeruginosa* | Aztreonam | - | - | - | - |
| *S. aureus* | Erythromycin | 2 | - | - | - |

The exact values in Table 2 were used to generate the proportions in Figure 2 of the main paper. Additional values are included here for combinations omitted.

#### C. Exact interpretability results

| Pathogen | Antimicrobial | AMR markers | IS element | rRNA | tRNA | Protein | No annotation |
|---|---|---|---|---|---|---|---|
| *A. baumannii* | Ceftazidime | 11 | 5 | – | 1 | 25 | 5 |
| | Ciprofloxacin | 13 | 1 | 1 | 2 | 71 | 2 |
| *E. coli* | Cefepime | 2 | – | – | – | – | – |
| | Cefotaxime | 3 | – | 6 | – | 4 | – |
| | Imipenem | 4 | 9 | 5 | 1 | 54 | 2 |
| *K. pneumoniae* | Meropenem | 7 | 1 | 2 | 3 | 13 | 1 |
| | Tobramycin | 2 | – | – | – | – | – |
| *P. aeruginosa* | Aztreonam | – | – | – | – | – | – |
| *S. aureus* | Erythromycin | 4 | 3 | – | – | 16 | 2 |

Table 3: Raw results of the interpretability study. Each important $k$-mer is mapped to a section of the reference genome that corresponds to one of the labels. All annotations were completed at the best performing $k$-mer length. *P. aeruginosa*–aztreonam is excluded as $k = 3$ is too small for reasonable annotation.

The values in Table 2 were the principal results of the interpretability study and were used to construct the figure of proportions present in the main paper. Models for all annotated combinations selected at least two $k$-mers that mapped to known AMR biomarkers.

#### D. Hyperparameter tuning ranges

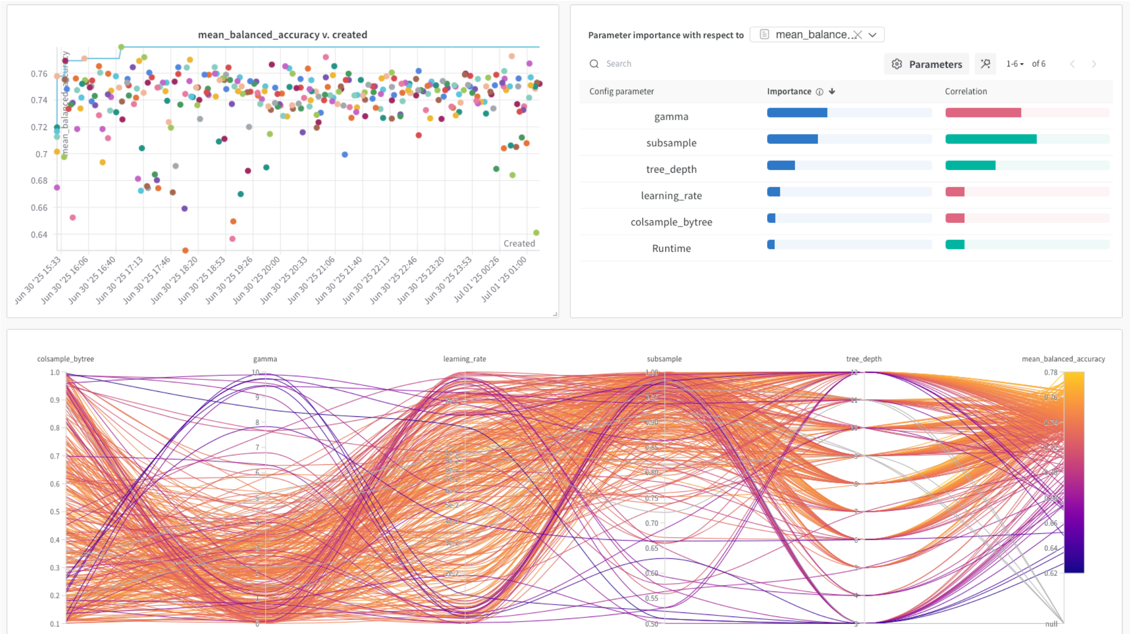


**Figure** 9: Results from XGBoost larger hyperparameter search

| Model | Parameter | Values | Sample method |
|---|---|---|---|
| XGBoost | | | |
| | `max_depth` | [3, 6, 9] | Selection |
| | `gamma` | 0.01 – 10.0 | Log uniform |
| | `subsample` | 0.6 – 1.0 | Uniform |
| LGR | | | |
| | `C` | 0.001 – 100 | Log uniform |
| | `penalty` | [`l1`, `elasticnet`] | Selection |
| | `solver` | saga | Fixed |
| | `max_iter` | [300, 400, 500] | Selection |

Table 4: Hyperparameters, search space, and sampling method for our XGBoost and Logistic Regression (LGR) models.

Hyperparameter ranges are presented in Table 4. We selected these based on empirical evidence and community knowledge of hyperparameters that commonly have a high impact on models of each type. Any unmentioned hyperparameters were left at their default values as set by the `scikit-learn` package. For XGBoost, we performed a larger sweep using the Weights and Biases platform with a total of five hyperparameters to evaluate the impact of each. Results from this sweep are shown in Figure 9. Based on the results, we eliminated `learning_rate` and `colsample_bytree` from our future searches.

#### E. Empirical time complexity of model types

Table 5: Time complexity for logistic regression on $k$-mer matrices, XGBoost on SVD-reduced matrices, and XGBoost on $k$-mer matrices. All values are in seconds, and ± indicate standard deviation across folds.

| **Pathogen** | **Antimicrobial** | **LGR** | **XGB/SVD** | **XGB** |
|---|---|---|---|---|
| *A. baumannii* | Ceftazidime | 29493.80 (± 14913.45) | 6839.31 (± 346.02) | 12383.77 (± 1500.87) |
| | Ciprofloxacin | 36074.68 (± 7605.13) | 5855.40 (± 308.20) | 17164.41 (± 3852.01) |
| *E. coli* | Cefepime | 44139.44 (± 27302.44) | 7459.06 (± 270.87) | 22219.30 (± 954.67) |
| | Cefotaxime | 16418.81 (± 5752.42) | 4024.35 (± 112.94) | 9043.18 (± 2065.35) |
| | Imipenem | 36281.97 (± 23158.08) | 5590.60 (± 295.28) | 25819.77 (± 794.19) |
| *K. pneumoniae* | Meropenem | 25083.16 (± 6503.89) | 3987.79 (± 82.35) | 8047.83 (± 1217.25) |
| | Tobramycin | 9251.05 (± 4174.75) | 3827.96 (± 123.12) | 9598.44 (± 6167.94) |
| *P. aeruginosa* | Aztreonam | 2.43 (± 0.07) | 17.59 (± 7.36) | 20.35 (± 5.17) |
| *S. aureus* | Erythromycin | 19106.03 (± 3668.04) | 3865.49 (± 207.04) | 5302.22 (± 365.96) |

We found that the additional processing required to compute the singular value decomposition still yielded lower overall time, since the XGBoost model had far fewer features to learn from. Logistic regression took much more time than XGBoost in most cases, except for extremely small datasets. XGBoost was also more consistent across folds in all cases except *P. aeruginosa*–aztreonam and *K. pneumoniae*–tobramycin, the two combinations where logistic regression also operated faster than XGBoost. These are also the two combinations with the fewest isolates among the set tested.

#### F. UCI dataset composition

Two of our nine species–antibiotic combinations were not included in the UCI dataset, and as such had to be omitted from the UCI evaluation. Additionally, *E. coli*–imipenem was omitted due to the extreme imbalance.

Table 6: UCI Dataset composition

| **Pathogen** | **Antimicrobial** | *n* | **Susceptible Samples** | **Resistant Samples** |
|---|---|---|---|---|
| *A. baumannii* | Ceftazidime | 39 | 32 (82%) | 7 (18%) |
| | Ciprofloxacin | 49 | 35 (71%) | 14 (29%) |
| *E. coli* | Cefepime | 2233 | 1725 (77%) | 508 (23%) |
| | Cefotaxime | 0 | - | - |
| | Imipenem | 2416 | 2410 (99.7%) | 6 (0.3%) |
| *K. pneumoniae* | Meropenem | 430 | 357 (83%) | 73 (17%) |
| | Tobramycin | 871 | 663 (76%) | 208 (24%) |
| *P. aeruginosa* | Aztreonam | 0 | - | - |
| *S. aureus* | Erythromycin | 541 | 320 (59%) | 221 (41%) |

MIC values were not reinterpreted, as they were already interpreted using CLSI M100 32nd edition by the original authors, which does not differ from the 35th edition for any of the seven relevant combinations.

#### G. Quality of phylogenetic folds

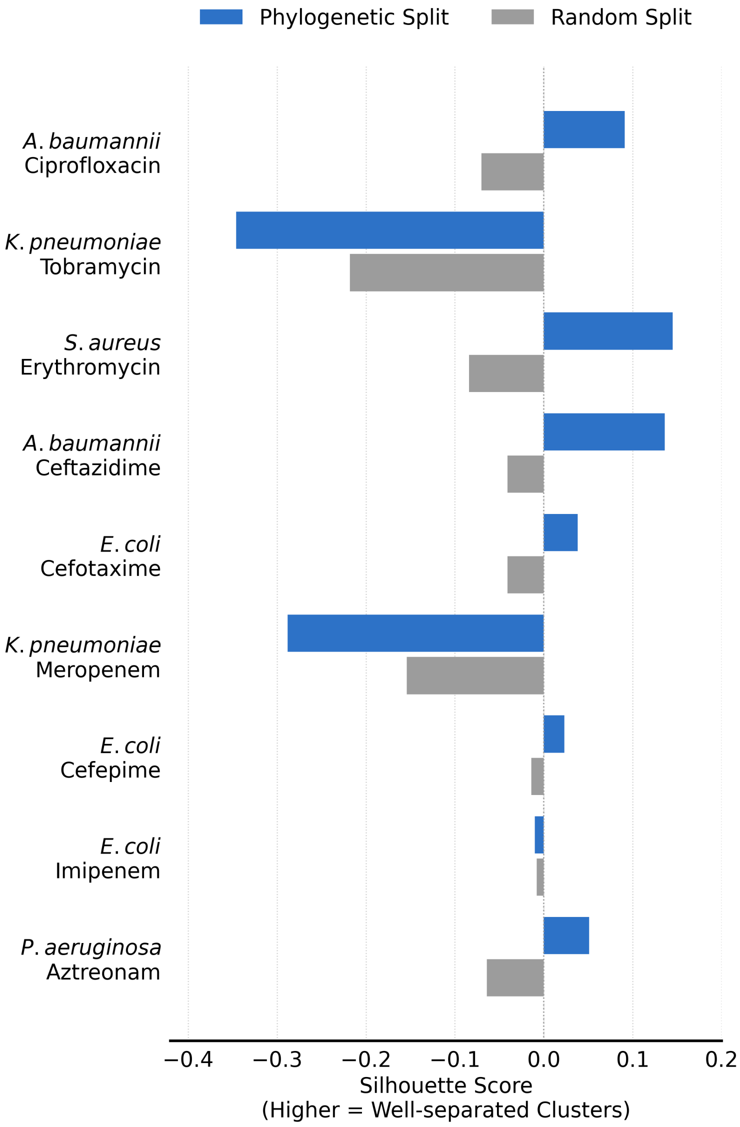


**Figure** 10: Silhouette scores depicting the separation between clusters obtained via random (grey) and phylogeny-aware (blue) splits of the dataset. Higher scores suggest that the clusters are well-separated. For six of nine combinations, phylogeny-aware splits preserve the distance between genomes of different clusters.

We first analyzed the silhouette scores of random and phylogeny-aware cross-validation folds to determine whether our phylogeny-aware splits create more well-separated groupings. Silhouette scores are between -1 and 1, where more negative values indicate that points from the nearest neighboring cluster are closer than points within a given cluster, on average. Higher scores indicate clusters that are more dissimilar from each other, or equivalently contain more similar genomes. Figure 10 shows that in six of nine cases, the phylogeny-aware cross-validation folds resulted in higher silhouette scores than random ones, indicating better separated clusters. This supports our claim that our phylogeny-aware folds correspond to train-test splits that have less in common. Interestingly, combinations with lower silhouette scores still saw a performance reduction when evaluating with phylogeny-aware folds over random folds.

#### H. SVD component selection

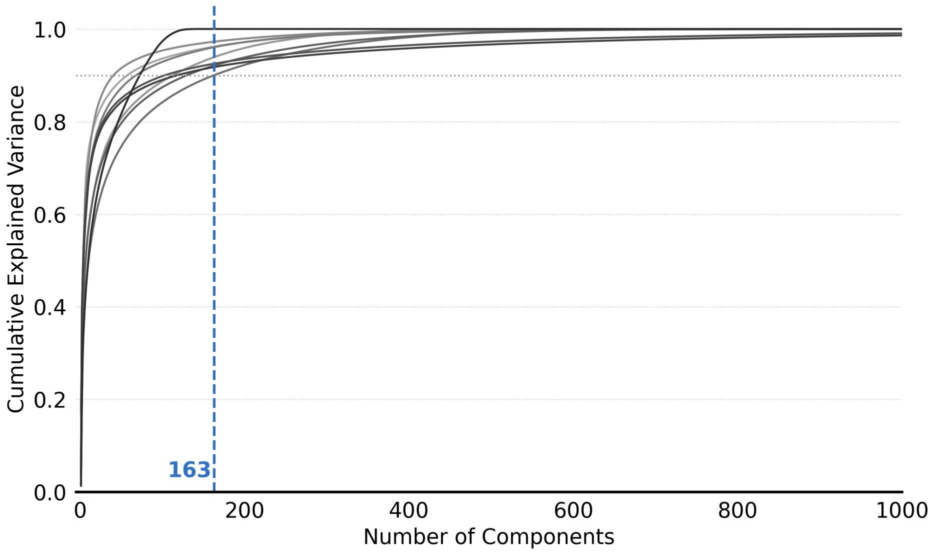


**Figure** 11: Cumulative explained variance ratio using truncated SVD with increasing numbers of components. The dashed blue line at 163 components indicates the threshold at which all combinations surpass 0.9.

Continuing the search for improved matrix efficiency, we experimented with the SVD to create extremely small and efficient feature matrices. During the tuning phase, we were primarily interested in determining the number of components required to achieve 0.9 explained variance. We selected this as a reasonable cut-off to capture most of the variation in the original data. We found 163 components to be enough to reach our 0.9 explained variance threshold across all combinations. After reaching this cut-off point, the increase in explained variance becomes very slow (Figure 11), indicating that there is little additional performance gain to be had.

### I. Hyperparameter search effects

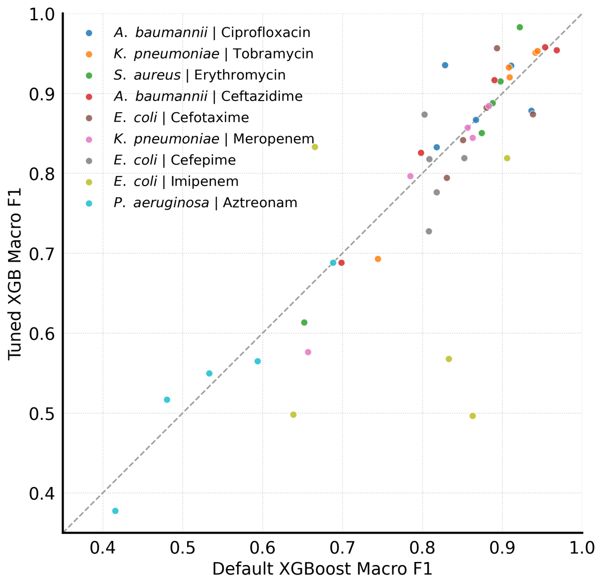


**Figure** 12: Macro F1 scores on each of the five phylogeny-aware folds for default and hyperparameter tuned XGBoost models. Folds are matched for both models.

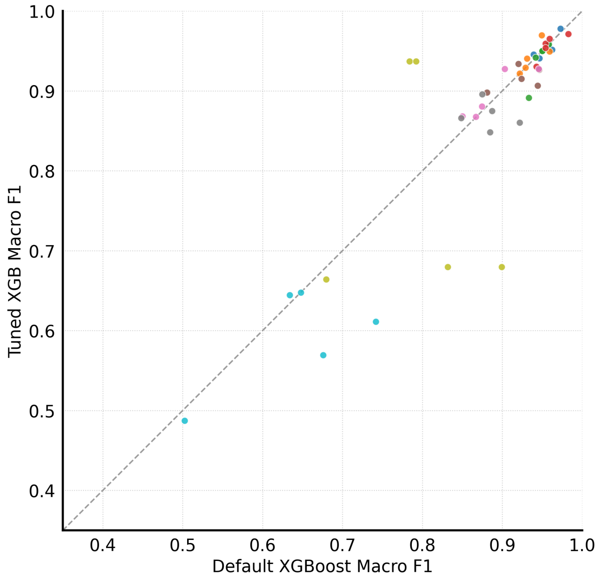


**Figure** 13: Random folds version of Figure 12.

We plotted the performance of the XGBoost models with default and tuned hyperparameters to ensure there were no systematic causes for the lack of performance increase. Visual inspection did not reveal any significant trends. Harder combinations, like *P. aeruginosa*–aztreonam and *E. coli*–imipenem, tended to have a larger standard deviation across default and tuned models. The majority of combinations are centered around y=x for both random and phylogeny-aware folds, verifying the Wilcoxon signed-rank test result.

### J. Additional metrics for key experiments

In AMR literature, major (misclassified as resistant) and very major (misclassified as susceptible) error rates are important metrics used to determine the clinical applicability of a system. Although we primarily relied on macro F1 scores for our analysis, we also tracked balanced accuracy, major error (ME) and very major error (VME) rates for all experiments. Here we provide values for a few of our key experiments: $k$-mer length (Table 7), model type (Table 8), and evaluation types (Table 9). These values are presented for comparison with other works in the field.

#### J.1. Evaluation Metrics

All reported metrics are averaged across the five cross-validation folds, except when reporting on an evaluation settings without folds (the UCI dataset and ResFinder baseline).

##### *J.1.1. Macro F1 Score*

For each class, precision and recall are defined as

$$\text{Precision}_c = \frac{TP_c}{TP_c + FP_c}, \tag{1}$$

$$\text{Recall}_c = \frac{TP_c}{TP_c + FN_c}, \tag{2}$$

where $TP_c$, $FP_c$, and $FN_c$ denote the numbers of true positives, false positives, and false negatives for class $c$, respectively.

The F1 score for class $c$ is

$$F_{1,c} = 2 \cdot \frac{\text{Precision}_c \times \text{Recall}_c}{\text{Precision}_c + \text{Recall}_c}. \tag{3}$$

For binary classification, the macro F1 score is computed as the unweighted mean of the class-specific F1 scores,

$$\text{Macro-}F_1 = \frac{1}{2}\left(F_{1,\text{susceptible}} + F_{1,\text{resistant}}\right). \tag{4}$$

For each experiment, the macro F1 score is computed independently on each of the five cross-validation test folds, and the reported value is the arithmetic mean across folds.

##### *J.1.2. Balanced Accuracy*

Balanced accuracy measures classification performance while accounting for class imbalance by averaging the recall of each class. For binary classification, it is defined as

$$\text{Balanced Accuracy} = \frac{1}{2}\left(\frac{TP}{TP + FN} + \frac{TN}{TN + FP}\right), \tag{5}$$

where resistant isolates are treated as the positive class.

##### *J.1.3. Major Error and Very Major Error*

Major error (ME) and very major error (VME) are standard evaluation metrics in antimicrobial susceptibility testing. A major error occurs when a susceptible isolate is incorrectly classified as resistant, whereas a very major error occurs when a resistant isolate is incorrectly classified as susceptible.

The major error rate is defined as

$$\text{ME} = \frac{FP}{TN + FP}, \tag{6}$$

representing the proportion of susceptible isolates misclassified as resistant.

Similarly, the very major error rate is

$$\text{VME} = \frac{FN}{TP + FN}, \tag{7}$$

representing the proportion of resistant isolates misclassified as susceptible.

#### J.2. Results

Table 7: Raw metrics for the four tested *k*-mer lengths across all nine combinations. Metrics include macro F1, balanced accuracy, major error (ME), and very major error (VME). All values are averaged across five folds, with standard deviations in brackets.

| Pathogen | Antimicrobial | Macro F1 | Bal. accuracy | ME | VME |
|---|---|---|---|---|---|
| | | | $k = 3$ | | |
| *A. baumannii* | Ceftazidime | 0.680 (± 0.107) | 0.699 (± 0.084) | 0.461 (± 0.153) | 0.140 (± 0.088) |
| | Ciprofloxacin | 0.710 (± 0.113) | 0.714 (± 0.108) | 0.459 (± 0.188) | 0.112 (± 0.106) |
| *E. coli* | Cefepime | 0.528 (± 0.037) | 0.531 (± 0.017) | 0.067 (± 0.085) | 0.871 (± 0.076) |
| | Cefotaxime | 0.522 (± 0.041) | 0.533 (± 0.034) | 0.798 (± 0.098) | 0.136 (± 0.037) |
| | Imipenem | 0.563 (± 0.103) | 0.543 (± 0.073) | 0.001 (± 0.001) | 0.912 (± 0.145) |
| *K. pneumoniae* | Meropenem | 0.556 (± 0.072) | 0.570 (± 0.069) | 0.492 (± 0.123) | 0.367 (± 0.046) |
| | Tobramycin | 0.603 (± 0.088) | 0.639 (± 0.074) | 0.461 (± 0.143) | 0.261 (± 0.082) |
| *P. aeruginosa* | Aztreonam | 0.539 (± 0.111) | 0.546 (± 0.123) | 0.618 (± 0.257) | 0.290 (± 0.161) |
| *S. aureus* | Erythromycin | 0.602 (± 0.086) | 0.611 (± 0.077) | 0.322 (± 0.153) | 0.456 (± 0.252) |
| | | | $k = 17$ | | |
| *A. baumannii* | Ceftazidime | 0.848 (± 0.118) | 0.841 (± 0.115) | 0.257 (± 0.167) | 0.061 (± 0.065) |
| | Ciprofloxacin | 0.890 (± 0.045) | 0.894 (± 0.053) | 0.171 (± 0.111) | 0.040 (± 0.017) |
| *E. coli* | Cefepime | 0.803 (± 0.055) | 0.773 (± 0.066) | 0.028 (± 0.037) | 0.427 (± 0.144) |
| | Cefotaxime | 0.870 (± 0.060) | 0.865 (± 0.068) | 0.223 (± 0.131) | 0.048 (± 0.027) |
| | Imipenem | 0.643 (± 0.170) | 0.618 (± 0.130) | 0.002 (± 0.004) | 0.761 (± 0.260) |
| *K. pneumoniae* | Meropenem | 0.792 (± 0.125) | 0.801 (± 0.092) | 0.148 (± 0.103) | 0.251 (± 0.157) |
| | Tobramycin | 0.888 (± 0.116) | 0.922 (± 0.044) | 0.060 (± 0.034) | 0.095 (± 0.066) |
| *P. aeruginosa* | Aztreonam | 0.438 (± 0.061) | 0.493 (± 0.066) | 0.604 (± 0.303) | 0.409 (± 0.289) |
| *S. aureus* | Erythromycin | 0.850 (± 0.141) | 0.852 (± 0.133) | 0.063 (± 0.063) | 0.233 (± 0.275) |
| | | | $k = 31$ | | |
| *A. baumannii* | Ceftazidime | 0.868 (± 0.114) | 0.872 (± 0.111) | 0.177 (± 0.141) | 0.078 (± 0.091) |
| | Ciprofloxacin | 0.830 (± 0.069) | 0.840 (± 0.065) | 0.239 (± 0.094) | 0.081 (± 0.064) |
| *E. coli* | Cefepime | 0.688 (± 0.076) | 0.671 (± 0.091) | 0.037 (± 0.044) | 0.622 (± 0.208) |
| | Cefotaxime | 0.739 (± 0.112) | 0.757 (± 0.109) | 0.388 (± 0.229) | 0.099 (± 0.042) |
| | Imipenem | 0.552 (± 0.060) | 0.538 (± 0.042) | 0.001 (± 0.003) | 0.923 (± 0.084) |
| *K. pneumoniae* | Meropenem | 0.720 (± 0.125) | 0.731 (± 0.088) | 0.221 (± 0.102) | 0.317 (± 0.174) |
| | Tobramycin | 0.885 (± 0.115) | 0.920 (± 0.044) | 0.060 (± 0.042) | 0.099 (± 0.065) |
| *P. aeruginosa* | Aztreonam | 0.455 (± 0.089) | 0.478 (± 0.065) | 0.581 (± 0.248) | 0.464 (± 0.210) |
| *S. aureus* | Erythromycin | 0.614 (± 0.139) | 0.661 (± 0.112) | 0.233 (± 0.222) | 0.446 (± 0.340) |
| | | | $k = 45$ | | |
| *A. baumannii* | Ceftazidime | 0.844 (± 0.132) | 0.845 (± 0.132) | 0.218 (± 0.167) | 0.093 (± 0.124) |
| | Ciprofloxacin | 0.809 (± 0.075) | 0.816 (± 0.054) | 0.280 (± 0.097) | 0.087 (± 0.084) |
| *E. coli* | Cefepime | 0.662 (± 0.112) | 0.654 (± 0.106) | 0.038 (± 0.054) | 0.654 (± 0.240) |
| | Cefotaxime | 0.760 (± 0.045) | 0.761 (± 0.041) | 0.381 (± 0.107) | 0.096 (± 0.043) |
| | Imipenem | 0.638 (± 0.134) | 0.637 (± 0.144) | 0.003 (± 0.005) | 0.723 (± 0.292) |
| *K. pneumoniae* | Meropenem | 0.717 (± 0.124) | 0.727 (± 0.095) | 0.252 (± 0.105) | 0.295 (± 0.160) |
| | Tobramycin | 0.890 (± 0.111) | 0.930 (± 0.027) | 0.042 (± 0.045) | 0.098 (± 0.072) |
| *P. aeruginosa* | Aztreonam | 0.538 (± 0.119) | 0.565 (± 0.143) | 0.583 (± 0.300) | 0.286 (± 0.140) |
| *S. aureus* | Erythromycin | 0.671 (± 0.102) | 0.692 (± 0.078) | 0.251 (± 0.220) | 0.364 (± 0.241) |

Table 8: Raw metrics for the three tested model types across all nine combinations. Metrics include macro F1, balanced accuracy, major error (ME), and very major error (VME). All values are averaged across five folds, with standard deviations in brackets.

| Pathogen | Antimicrobial | Macro F1 | Bal. accuracy | ME | VME |
|---|---|---|---|---|---|
| | | **Logistic Regression with 1M *k*-mer matrices** | | | |
| *A. baumannii* | Ceftazidime | 0.840 (± 0.118) | 0.842 (± 0.113) | 0.204 (± 0.140) | 0.112 (± 0.124) |
| | Ciprofloxacin | 0.730 (± 0.305) | 0.782 (± 0.192) | 0.204 (± 0.163) | 0.232 (± 0.409) |
| *E. coli* | Cefepime | 0.660 (± 0.082) | 0.647 (± 0.078) | 0.045 (± 0.067) | 0.660 (± 0.202) |
| | Cefotaxime | 0.786 (± 0.110) | 0.791 (± 0.114) | 0.338 (± 0.226) | 0.080 (± 0.050) |
| | Imipenem | 0.570 (± 0.112) | 0.582 (± 0.117) | 0.003 (± 0.005) | 0.833 (± 0.236) |
| *K. pneumoniae* | Meropenem | 0.725 (± 0.091) | 0.733 (± 0.061) | 0.260 (± 0.104) | 0.274 (± 0.153) |
| | Tobramycin | 0.864 (± 0.122) | 0.912 (± 0.034) | 0.067 (± 0.053) | 0.110 (± 0.093) |
| *P. aeruginosa* | Aztreonam | 0.439 (± 0.047) | 0.479 (± 0.078) | 0.827 (± 0.205) | 0.215 (± 0.191) |
| *S. aureus* | Erythromycin | 0.630 (± 0.140) | 0.668 (± 0.129) | 0.146 (± 0.175) | 0.518 (± 0.260) |
| | | **XGBoost with SVD-reduced *k*-mer matrices** | | | |
| *A. baumannii* | Ceftazidime | 0.720 (± 0.182) | 0.788 (± 0.113) | 0.119 (± 0.086) | 0.305 (± 0.266) |
| | Ciprofloxacin | 0.687 (± 0.195) | 0.761 (± 0.140) | 0.188 (± 0.094) | 0.290 (± 0.270) |
| *E. coli* | Cefepime | 0.638 (± 0.058) | 0.617 (± 0.040) | 0.018 (± 0.010) | 0.749 (± 0.079) |
| | Cefotaxime | 0.724 (± 0.094) | 0.698 (± 0.085) | 0.584 (± 0.176) | 0.020 (± 0.010) |
| | Imipenem | 0.557 (± 0.090) | 0.561 (± 0.108) | 0.001 (± 0.001) | 0.878 (± 0.217) |
| *K. pneumoniae* | Meropenem | 0.642 (± 0.122) | 0.657 (± 0.083) | 0.345 (± 0.092) | 0.342 (± 0.205) |
| | Tobramycin | 0.789 (± 0.048) | 0.801 (± 0.072) | 0.223 (± 0.096) | 0.174 (± 0.078) |
| *P. aeruginosa* | Aztreonam | 0.458 (± 0.102) | 0.472 (± 0.109) | 0.724 (± 0.197) | 0.333 (± 0.158) |
| *S. aureus* | Erythromycin | 0.536 (± 0.174) | 0.615 (± 0.103) | 0.152 (± 0.178) | 0.618 (± 0.337) |
| | | **XGBoost with 1M *k*-mer matrices** | | | |
| *A. baumannii* | Ceftazidime | 0.868 (± 0.114) | 0.872 (± 0.111) | 0.177 (± 0.141) | 0.078 (± 0.091) |
| | Ciprofloxacin | 0.890 (± 0.045) | 0.894 (± 0.053) | 0.171 (± 0.111) | 0.040 (± 0.017) |
| *E. coli* | Cefepime | 0.803 (± 0.055) | 0.773 (± 0.066) | 0.028 (± 0.037) | 0.427 (± 0.144) |
| | Cefotaxime | 0.870 (± 0.060) | 0.865 (± 0.068) | 0.223 (± 0.131) | 0.048 (± 0.027) |
| | Imipenem | 0.643 (± 0.170) | 0.618 (± 0.130) | 0.002 (± 0.004) | 0.761 (± 0.260) |
| *K. pneumoniae* | Meropenem | 0.792 (± 0.125) | 0.801 (± 0.092) | 0.148 (± 0.103) | 0.251 (± 0.157) |
| | Tobramycin | 0.890 (± 0.111) | 0.930 (± 0.027) | 0.042 (± 0.045) | 0.098 (± 0.072) |
| *P. aeruginosa* | Aztreonam | 0.539 (± 0.111) | 0.546 (± 0.123) | 0.618 (± 0.257) | 0.290 (± 0.161) |
| *S. aureus* | Erythromycin | 0.850 (± 0.141) | 0.852 (± 0.133) | 0.063 (± 0.063) | 0.233 (± 0.275) |

Table 9: Raw metrics for the three tested evaluation settings across all nine combinations. Metrics include macro F1, balanced accuracy, major error (ME), and very major error (VME). Cross-validation results are averaged across five folds, with standard deviations in brackets. UCI dataset results are reported based on classification of the entire dataset, and so do not have standard deviations.

| Pathogen | Antimicrobial | Macro F1 | Bal. accuracy | ME | VME |
|---|---|---|---|---|---|
| | | **Phylogeny-aware CV** | | | |
| *A. baumannii* | Ceftazidime | 0.868 (± 0.114) | 0.872 (± 0.111) | 0.177 (± 0.141) | 0.078 (± 0.091) |
| | Ciprofloxacin | 0.890 (± 0.045) | 0.894 (± 0.053) | 0.171 (± 0.111) | 0.040 (± 0.017) |
| *E. coli* | Cefepime | 0.803 (± 0.055) | 0.773 (± 0.066) | 0.028 (± 0.037) | 0.427 (± 0.144) |
| | Cefotaxime | 0.870 (± 0.060) | 0.865 (± 0.068) | 0.223 (± 0.131) | 0.048 (± 0.027) |
| | Imipenem | 0.643 (± 0.170) | 0.618 (± 0.130) | 0.002 (± 0.004) | 0.761 (± 0.260) |
| *K. pneumoniae* | Meropenem | 0.792 (± 0.125) | 0.801 (± 0.092) | 0.148 (± 0.103) | 0.251 (± 0.157) |
| | Tobramycin | 0.890 (± 0.111) | 0.930 (± 0.027) | 0.042 (± 0.045) | 0.098 (± 0.072) |
| *P. aeruginosa* | Aztreonam | 0.539 (± 0.111) | 0.546 (± 0.123) | 0.618 (± 0.257) | 0.290 (± 0.161) |
| *S. aureus* | Erythromycin | 0.850 (± 0.141) | 0.852 (± 0.133) | 0.063 (± 0.063) | 0.233 (± 0.275) |
| | | **Random CV** | | | |
| *A. baumannii* | Ceftazidime | 0.956 (± 0.016) | 0.957 (± 0.017) | 0.063 (± 0.027) | 0.023 (± 0.007) |
| | Ciprofloxacin | 0.955 (± 0.014) | 0.958 (± 0.015) | 0.063 (± 0.026) | 0.020 (± 0.007) |
| *E. coli* | Cefepime | 0.869 (± 0.018) | 0.854 (± 0.026) | 0.026 (± 0.007) | 0.267 (± 0.054) |
| | Cefotaxime | 0.916 (± 0.014) | 0.906 (± 0.013) | 0.159 (± 0.022) | 0.029 (± 0.012) |
| | Imipenem | 0.779 (± 0.144) | 0.722 (± 0.152) | 0.000 (± 0.001) | 0.556 (± 0.304) |
| *K. pneumoniae* | Meropenem | 0.894 (± 0.031) | 0.896 (± 0.029) | 0.097 (± 0.017) | 0.112 (± 0.049) |
| | Tobramycin | 0.942 (± 0.019) | 0.944 (± 0.016) | 0.060 (± 0.029) | 0.053 (± 0.036) |
| *P. aeruginosa* | Aztreonam | 0.592 (± 0.067) | 0.593 (± 0.067) | 0.536 (± 0.113) | 0.278 (± 0.088) |
| *S. aureus* | Erythromycin | 0.938 (± 0.027) | 0.938 (± 0.027) | 0.046 (± 0.021) | 0.078 (± 0.043) |
| | | **UCI** | | | |
| *A. baumannii* | Ceftazidime | 1.000 | 1.000 | 0.000 | 0.000 |
| | Ciprofloxacin | 0.976 | 0.980 | 0.020 | 0.000 |
| *E. coli* | Cefepime | 0.856 | 0.897 | 0.058 | 0.045 |
| | Cefotaxime | – | – | – | – |
| | Imipenem | – | – | – | – |
| *K. pneumoniae* | Meropenem | 0.878 | 0.921 | 0.072 | 0.007 |
| | Tobramycin | 0.975 | 0.982 | 0.007 | 0.011 |
| *P. aeruginosa* | Aztreonam | – | – | – | – |
| *S. aureus* | Erythromycin | 0.942 | 0.945 | 0.015 | 0.041 |
